# Differential gene expression underlying the biosynthesis of Dufour’s gland signals in *Bombus impatiens*

**DOI:** 10.1101/2022.04.12.488013

**Authors:** Nathan Derstine, David Galbraith, Gabriel Villar, Etya Amsalem

## Abstract

Pheromones regulating social behavior are one of the most explored phenomena in social insects. However, compound identity, biosynthesis and their genetic regulation are known in only a handful of species. Here we conducted chemical and RNA-seq analyses of the Dufour’s gland in the bumble bee *Bombus impatiens* and examined the signals and the pathways regulating signal production in queens and workers. Across Hymenopterans, the Dufour’s gland contains mostly long-chained hydrocarbons and esters that signal reproductive and social status in several bee species. In bumble bees, the Dufour’s gland contains queen- and worker-specific esters, in addition to terpenes and terpene-esters only found in gynes and queens. These compounds are assumed to be synthesized de novo in the gland, however, their genetic regulation is unknown. A whole transcriptome gene expression analysis of the gland in queens, gynes, queenless and queenright workers showed distinct transcriptomic profiles, with thousands of differentially expressed genes between the groups. Workers and queens express genes associated with key enzymes in the biosynthesis of wax esters, while queens and gynes preferentially express key genes in terpene biosynthesis. In contrast, no genes were differentially expressed in queenless and queenright workers, despite differences in their Dufour’s gland chemistry and reproductive state, suggesting the quantitative differences in worker secretion are not regulated at the level of production. Overall, our data demonstrate gland-specific regulation of chemical signals associated with social behavior and identifies genes and pathways regulating caste-specific chemical signals in social insects.

## Introduction

Reproductive division of labor between a primary egg-layer and non-reproductive females is a defining feature of eusocial insects (Wilson, 1971), and is often maintained using chemical signals produced in exocrine glands (Amsalem, 2020; A. Hefetz, 2019). Such signals have been identified in numerous insect species (Hoover, Keeling, Winston, & Slessor, 2003; Steiger & Stokl, 2018; Steitz & Ayasse, 2020), and while they can be quite diverse (Amsalem, 2020; A. Hefetz, 2019), they are commonly derivatives of fatty acid biosynthesis (Yew & Chung, 2015). Fatty acid derivatives such as hydrocarbons, wax esters, fatty alcohols and acids are frequently represented as signal molecules, and thus may share elements related to their biosynthesis and genetic regulation. In contrast, terpenes, another frequently represented class of signal molecules, derive from a separate pathway (Bellés, Martín, & Piulachs, 2005). Despite the importance of chemical signals for regulating reproduction and social behavior, their molecular and biosynthetic underpinnings are often poorly understood. Better understanding of signal biosynthesis and its genetic regulation enables direct manipulations of key genes to examine mechanisms of signal production (Yan & Liebig, 2021), and is a crucial step towards understanding the evolution of signals and sociality.

Signals that are derivatives of fatty acid biosynthesis are common across social insect species. For instance, hydrocarbons serve primely to prevent insect desiccation (Menzel et al., 2019), but evolved as signals reflecting species, caste, age, task, reproductive and social status in many species (Blomquist & Bagnères, 2010; Derstine, Gries, Zhai, Jimenez, & Gries, 2018; Dietemann, Peeters, Liebig, Thivet, & Holldobler, 2003; Endler et al., 2004; Oi et al., 2015; Smith, Holldober, & Liebig, 2009; Smith, Millar, Hanks, & Suarez, 2013). Likewise, various wax esters comprise the brood pheromone of honey bees (Yves Le Conte, Mohammedi, & Robinson, 2001), and play a role in the reproductive signaling of worker bumble bees (Amsalem, Twele, Francke, & Hefetz, 2009), and macrocyclic lactones (cyclic esters) serve as queen pheromones in halictid bees (Steitz & Ayasse, 2020; Steitz, Brandt, Biefel, Minat, & Ayasse, 2019). Furthermore, the main component of the honey bee queen mandibular pheromone is a fatty acid derived hydroxy-acid which regulates worker reproduction (Erika Plettner, Slessor, & Winston, 1998; Slessor, Kaminski, King, & Winston, 1990). The common biosynthetic origin of these molecules and their role in regulating reproduction suggests an underlying genetic link.

The Dufour’s gland (DG) is an exocrine gland which produces a range of chemical signals that regulate disparate aspects of social life in Hymenopterans (Mitra, 2013), providing an excellent opportunity to study the molecular determinants of chemical signals and their biosynthesis. The gland is specific to females and is located close to the sting apparatus where it opens into the oviduct. Depending on the species, it contains various compounds such as hydrocarbons, esters, lactones, terpenes and terpene esters, and fatty acids, some of which regulate different social functions. For instance, DG terpene esters in *Gnamptogenys striatula* serve as trail pheromones (Blatrix, Schulz, Jaisson, Francke, & Hefetz, 2002), and macrocyclic lactones in the social Halictid *Lasioglossum malachurum*, serve as a queen pheromone (Steitz & Ayasse, 2020). In bumble bees and honey bees, worker and queen specific esters in the DG act as sterility and fertility signals, respectively, advertising female fecundity (Amsalem, Shpigler, Bloch, & Hefetz, 2013; Amsalem et al., 2009; Dor, Katzav-Gozansky, & Hefetz, 2005). In honey bees, it has been shown that the esters are produced de-novo in the gland (Katzav-Gozansky, Soroker, & Hefetz, 2000) while the hydrocarbons are synthesized in oenocytes and transported into the gland via lipophorins (Soroker & Hefetz, 2000), indicating that the gland is not only a reservoir for the signals but also the place where the active components of the signal are being produced.

Bumble bees provide an excellent model species to examine the biosynthesis of reproductive signaling. The variation found between castes and reproductive states in the DG secretion, and the ability to replicate this variation by manipulating the social environment and mating status facilitates comparative experiments. Furthermore, the composition of the DG has been extensively studied in several bumble bee species, and contains saturated and unsaturated hydrocarbons, wax and acetate esters, fatty acids, and terpenes in blends unique to both species and caste (Cahlíková, Hovorka, Ptáček, & Valterová, 2004; Derstine et al., 2021; Tengö et al., 1991). These compounds vary with age and also within workers of different reproductive state or social condition (Derstine et al., 2021; Urbanová, Cahlíková, Hovorka, Ptáček, & Valterová, 2008). In *B. terrestris*, wax esters in the DG disappear in workers as their ovaries develop and it has been suggested that their presence signals sterility (Amsalem et al., 2009), reducing unnecessary aggression as dominance hierarchies are established (Amsalem & Hefetz, 2010) and/or advertising the risky work of forager bees (Amsalem et al., 2013). In *B. impatiens*, worker-specific wax esters correlate with age and ovarian activation, while queens and gynes (young, non-reproductive queens) produce terpenes and terpene esters not found in workers (Derstine et al., 2021). Similar terpenes are frequently found in the labial glands of *Bombus sp*. males and serve as sex pheromones (Valterová, Martinet, Michez, Rasmont, & Brasero, 2019). This existing information makes bumble bees an excellent model to study the genetic regulation underlying the production of these signals.

Despite the importance of DG esters and terpenes as signal molecules, little is known of the genes underlying their biosynthesis. Wax esters are formed from the condensation of fatty acids and alcohols. Their synthesis involves the elongation of fatty acids, conversion of fatty acyl-CoAs to fatty alcohol, and final esterification step. Two enzymes are likely involved in the esterification: a fatty acyl reductase (FAR), and a wax synthase (WS) which is a specialized acyltransferase (Teerawanichpan & Qiu, 2010; Wenning, Yu, David, Nielsen, & Siewers, 2017). FAR enzymes are responsible for the production of primary fatty alcohols, which can serve as signals themselves and also as precursors to a variety of esters (Matoušková et al., 2008; Valterová et al., 2019). Insects have multiple FAR enzymes with varying levels of substrate specificity, and FAR enzymes in hymenopterans have undergone an expansion implicated in shaping species-specific pheromone communication (Tupec et al., 2019). While current evidence suggests that a WS catalyzes the esterification of a fatty acyl-CoA and a primary fatty alcohol to produce a wax ester, genes with wax synthase activity have only been identified in non-insect organisms, such as mammals, plants, yeasts, and bacteria, (Yang et al., 2012). In many organisms, a multi-functional gene in the acyltransferase family (such as diacylglycerol acyltransferase or DGAT) catalyzes the production of wax esters, but also mono, di, or triglycerides (Kalscheuer & Steinbüchel, 2003; F. Li et al., 2008; Teerawanichpan & Qiu, 2010; Wenning et al., 2017; Yen, Monetti, Burri, & Farese Jr, 2005). In *B. impatiens*, worker specific wax esters are formed from acid and alcohol components less than 16 carbons long, as opposed to the queen-specific wax esters which are longer, indicating a chain-shortening beta-oxidation process in workers, but not in queens, prior to esterification. Overall, the caste specificity of the wax esters in *B. impatiens* Dufour’s gland likely results from differences in beta-oxidation activity and the expression of FAR enzymes, which supply primary alcohols of varying lengths to be esterified by a relatively non-specific wax synthase/DGAT.

Terpenes produced by the queen caste are synthesized via isoprenoid pathway and share a biosynthetic origin with the terpenes serving as sex pheromones in the labial glands of *Bombus males* (Brabcova et al., 2015; Prchalova et al., 2016; Valterová et al., 2019). This pathway gives rise to an enormous diversity of compounds including monoterpenes, sesquiterpenes, and juvenile hormones. These compounds function in defense, and act as chemical signals mediating myriad behaviors across many taxa. For instance, dihydrofarnesol and other terpenoids mediate sexual attraction in bumble bees (Valterová et al., 2019), and a sesquiterpene blend mediates aggregation and mating in a chrysomelid beetle (Beran, Jiménez-Alemán, et al., 2016) and in certain stink bugs (Zahn, Moreira, & Millar, 2008). Furthermore, fungus gnats employ sesquiterpene sex pheromone (Andreadis, Cloonan, Myrick, Chen, & Baker, 2015), the monoterpene β-ocimene acts as a brood pheromone in honey bees (Maisonnasse, Lenoir, Beslay, Crauser, & Le Conte, 2010) and as an anti-aphrodisiac pheromone in *Heliconius* butterflies (Darragh et al., 2021), while the terpene ester methyl geranate does the same in burying beetles (Engel et al., 2016). Beyond their role as interindividual signals, terpenes regulate physiological processes as hormone production, electron transport and cellular signaling (Bellés et al., 2005; Morgan, 2010).

To examine the biosynthesis and genetic regulation of wax esters and terpenes in the Dufour’s gland of *B. impatiens* females, we analyzed the chemical composition of the gland and conducted RNAseq analyses of the gland in queen, gynes and workers (both queenright and queenless). We examine variation in gene expression associated with caste, mating status, and fertility. We hypothesized that: 1) genes in the fatty acid biosynthetic pathway related to chain shortening and beta-oxidation would be upregulated in workers groups which produce shorter wax esters than queens, and would be highest in queenright workers, which produce more of these compounds compared to queenless workers, 2) FAR and acyltransferase genes involved in ester formation would be downregulated in queens that produce little to no wax esters, and 3) genes in the isoprenoid biosynthetic pathway would be upregulated in gynes which produce the most terpenes compared to queens and workers.

## Methods

### Insect Rearing and Sampling

Bees were collected from *B. impatiens* colonies that were obtained from Koppert Biological Systems (Howell, Michigan, USA). Colonies were maintained in nest-boxes at a constant darkness, temperature of 28–30°C and 60% relative humidity and supplied *ad libitum* with 60% sugar solution and honeybee-collected pollen (purchased from Koppert). These colonies were used as a source for mated, egg-laying queens (hereafter “queens”), newly-emerged, unmated queens (hereafter “gynes”), and newly-emerged workers. Upon emergence, workers were assigned to one of two treatments: queenright workers (QRW) and queenless and brood-less workers (QLW). QRW were individually labeled and remained in their natal colony in the presence of the queen, brood, and nestmates until collection at the desired age, while QLW were housed in a plastic cage (11 cm diameter × 7 cm height) in groups of 3-6 workers until collection. Both groups of workers were collected at the age of 7 days. Since the presence of brood can impact worker reproduction (Starkey, Brown, & Amsalem, 2019), QLW cages were monitored daily for egg laying, and newly-laid eggs were removed. Gynes were collected upon emergence from late-season colonies and housed individually in small cages until sampling on days 3-4. Queens were sampled from colonies with ongoing egg-laying and worker production, indicating that they were successfully mated, and were > 8 weeks old from the emergence of the first worker. The time points of the treatment groups were selected to capture differences in phenotype of both ovarian activation and the DG secretion based on previous work (Derstine et al., 2021).

These experimental procedures were repeated twice; once to generate samples for chemical analysis of the DG, and a second time to generating samples for RNA extraction and subsequent RNAseq analysis. For chemical analyses, QRW (n = 15), QLW (n = 15), and gynes (n = 20) were sampled from a total of 8 colonies (minimum 4 bees per colony). Queens (n = 20) each originated from a different colony. RNA was extracted from pools of 5-10 workers (58 QRW and 48 QLW), 3-7 gynes (n=33) and single queens (n=6) and collected from 15 colonies resulting in 24 samples (6 QLW, 6 QRW, 6 gynes and 6 queens). Detailed information on the number of glands used per sample and their colony of origin is found in Table S1.

### Ovarian activation

Oocyte size was measured in all samples. Ovaries were removed and placed in a drop of distilled water, and the largest three terminal oocytes across both ovaries (at least one from each ovary) were measured with an eyepiece micrometer (Amsalem & Hefetz, 2010). The average terminal oocyte was used as a measure of ovary activation for each bee.

### Chemical analysis

Bees were dissected under a stereomicroscope in distilled water and the gland was removed. Glands were extracted for 24 hours in 150 µL hexane containing internal standard of eicosane (C_20_) at 20ng/µL, before being analyzed by gas-chromatography flame ionization detection (GC-FID), and GC-mass spectrometry.

DG components of *B. impatiens* females were identified in a previous study (Derstine et al., 2021). To quantify the components in this study, DG extracts were analyzed on a Trace 1310 GC (Thermo Fisher, Waltham, MA USA) equipped with a flame-ionization detector (FID) and a TG-5MS column (0.25 mm id x 30 m × 0.25 µm film thickness; Thermo Fisher). The oven temperature was programmed at 60°C for 1 min, increased to 120°C at 15°C/min, then increased to 300°C at 4°C/min and held at 300°C for 5 min. The injector port and FID were held constant at 250°C and 320°C, respectively.

### RNA extraction

DGs were dissected by removing the whole sting complex of a dry-ice anesthetized bee and placing it in a drop of RNAlater (Invitrogen, Carlsbad, CA) to keep the tissue cold and prevent degradation. Individual DGs were removed from the sting complex using clean forceps and pooled according to treatment group in sterilized 2 mL FastPrep tubes containing six 2 mm zirconia beads and 250 µL Trizol (Invitrogen, Carlsbad, CA). Individual DGs were pooled to obtain sufficient amounts of RNA for downstream analyses. Worker samples contained RNA from 5-10 glands, gynes from 5-7 glands, and queens from a single gland (Table S1). Samples were frozen at -80° C and stored until processing. Total RNA was isolated using a PicoPure kit according to the manufacturer instructions. RNA quantity was measured using a NanoDrop One™.

### RNA-sequencing and differential gene expression analysis

Twenty-four cDNA libraries were prepared using Illumina TruSeq protocols (n = 6 per treatment in QLW, QRW, gyne, and queen groups). RNA-sequencing was performed using single end 75nt read lengths at the Penn State Genomics Core Facility on an Illumina NextSeq system. FastQC (Andrews, 2010) and MultiQC (Ewels, Magnusson, Lundin, & Kaller, 2016) were used to identify potential sequencing issues such as low quality reads. According to the recommended usage of Trimmomatic (Bolger, Lohse, & Usadel, 2014), reads with a Phred score below 25 and length below 36 bp were removed. The reads were then processed with Trimmomatic v0.039 to remove TruSeq3-SE adapter sequences and to trim low-quality bases from the ends of the reads. The processed reads were aligned to the *Bombus impatiens* genome assembly GCF_000188095.2 (Sadd et al., 2015) using STAR v2.7 (Dobin et al., 2013) and read counts were calculated using the standard RSEM pipeline (B. Li & Dewey, 2011), producing count data for 10633 genes, with an average of 18,309,015 reads per sample.

Differential gene expression analyses were conducted using the packages edgeR (Robinson, McCarthy, & Smyth, 2010) and limma (Law et al., 2016; Ritchie et al., 2015) in R v. 4.1.3. First, lowly expressed genes (1064 which had a sum of transcript counts < 20 across all samples) were removed from analyses. The *voom* method (Law, Chen, Shi, & Smyth, 2014) was used to estimate the mean-variance relationship of log transcript counts per million (logCPM) and create a precision weight for every normalized observation. These weights are used when fitting linear models to the gene expression data to account for variable standard deviation inherent in comparing count data across a large scale. Differentially expressed genes (DEGs) were identified between specific contrasts of treatment groups (queen vs. gyne, queen vs. QRW, queen vs. QLW, gyne vs. QRW, gyne vs. QLW, queens vs. workers), and multiple testing was controlled using Benjamini and Hochberg’s false discovery rate at 0.05 to adjust P-value cutoffs (Benjamini & Hochberg, 1995). We analyzed the contrasts of QLW vs. QRW to examine the effect of reproductive status and social condition on worker gene expression, queen vs. gyne to examine the same effects in queens, and queen vs QLW to examine the effect of caste on gene expression in reproductively active bees. T-distributed stochastic neighbor embedding (t-SNE) was used to visualize sample clustering based on the full gene count matrix with the Rtsne and ggplot2 packages (Krijthe & Van der Maaten, 2015; Wickham, 2011). A heatmap of selected DEGs was created using the heatmap.2 function from the gplots package (Warnes, Bolker, Bonebakker, Gentleman, & Huber, 2016). An upset plot of DEG sets was created using the UpSetR package (Conway, Lex, & Gehlenborg, 2017). Volcano plots of selected contrasts were made using the EnhancedVolcano package (Blighe, Rana, & Lewis, 2021).

### Statistical analyses

Statistical analyses of DG compounds were performed using R (version 4.0.5) in RStudio (version 1.2.5033). To examine chemical differences in the DG secretion between females of different caste (queens, workers) and social conditions (QRW, QLW), permutational analysis of variance (PERMANOVA) was performed using the *adonis* function from the vegan package (Oksanen et al., 2013) on relative peak areas, with social condition and caste as grouping variables. Pairwise comparisons between groups was performed with *pairwise*.*perm*.*manova* function from the RVAideMemoire package (Hervé & Hervé, 2020) using the Wilks test and FDR correction for multiple testing. Only peaks > 0.5% of the total peak area in at least one group were included. Non-metric multidimensional scaling plots (NMDS) using esters, alkanes, alkenes, or all compounds were used to visualize the results.

Mean terminal oocyte size was compared between treatment groups using a one-way ANOVA followed by Tukey’s HSD post-hoc test. When comparing the percentage of chemical classes between treatment groups, the data were not always normally distributed (Shapiro-Wilks test > 0.05) or had unequal variances (Levene’s test > 0.05), and thus relative and absolute amounts of chemical classes were compared between treatment groups using the non-parametric Kruskal-Wallis test. Dunn’s post-hoc test from the dunn.test package (Dinno & Dinno, 2017) and Benjamini and Hochberg multiple comparison correction were performed to provide contrasts between treatment groups.

## Results

### Social condition and reproductive phenotype

Ovarian activation was significantly different between treatment groups (One-way ANOVA, F_3,210_= 797.8, p < 0.001). Queens and queenless workers (QLW) displayed higher ovary activation compared to gynes and queenright workers (QRW) (Fig 1a). On average, queens had significantly larger terminal oocytes than gynes (3.4 ± 0.03 and 0.29 ± 0.03, mean mm ± SE, respectively, ANOVA followed by Tukey HSD post-hoc test, p < 0.001), and QLW had significantly larger terminal oocytes than QRW (2.56 ± 0.07 and 0.41 ± 0.04, mean mm ± SE, respectively, ANOVA followed by Tukey HSD post-hoc test, p < 0.001). In groups with activated ovaries, queen terminal oocyte size was also larger than QLW (ANOVA followed by Tukey HSD post-hoc test, p < 0.001). Gynes and QRW ovaries were not significantly different.

**Figure 1.**
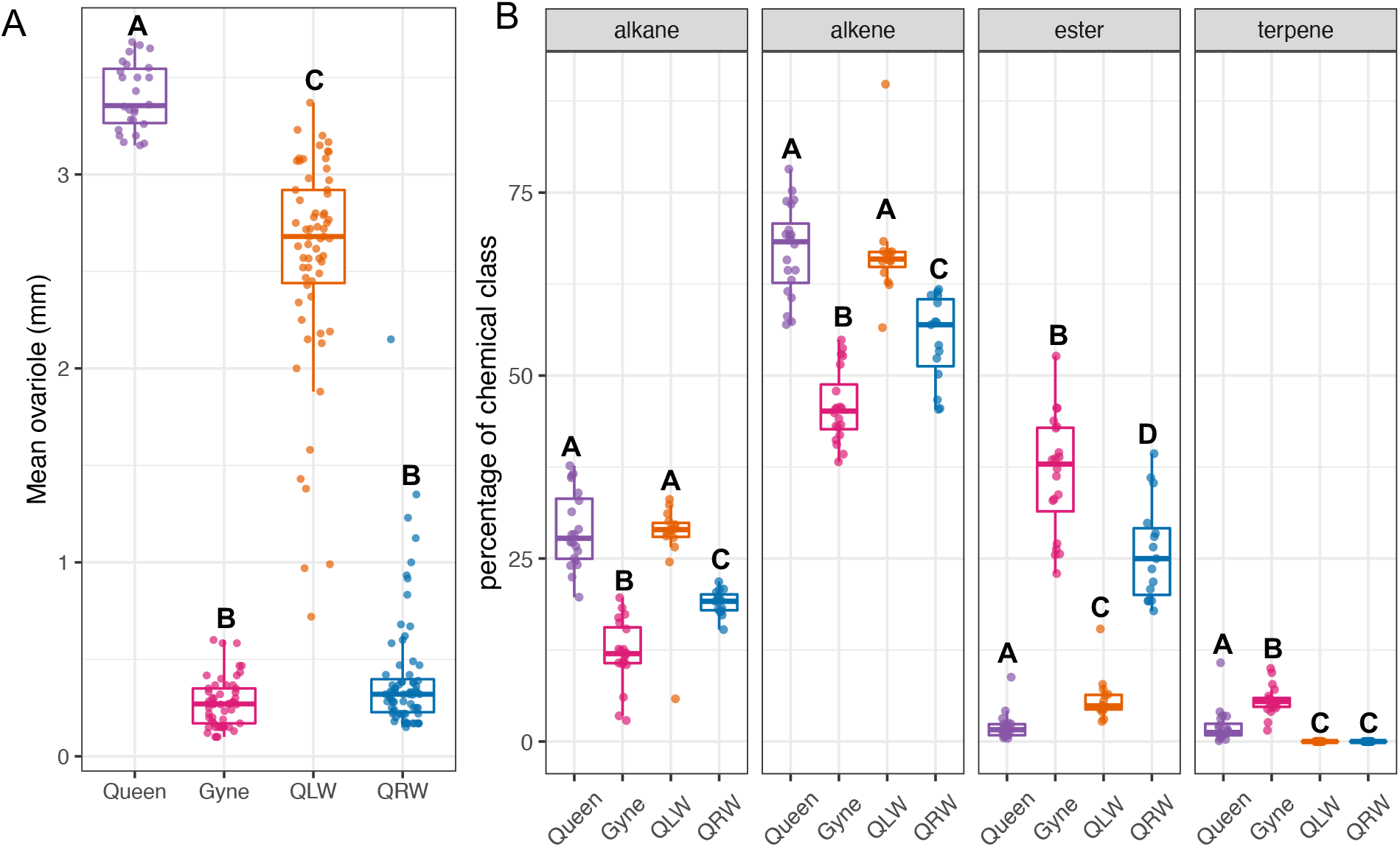
A - Average terminal oocyte length of bees used in chemical and RNAseq in four groups of *B. impatiens* females: 7-days-old queenright workers (QRW, n=73), 7-days-old queenless workers (n=63), 3-4 days old gynes (n=53) and active queens (n=26). B – Percentages of specific chemical classes of the *B. impatiens* DG secretion in four groups of *B. impatiens* females: 7-days-old queenright workers (QRW, n=15), 7-days-old queenless workers (n=15), 3-4 days old gynes (n=20) and active queens (n=20). Different letters denote significant differences at p < 0.05 (mean ovariole - One-way ANOVA with Tukey HSD post-hoc test; percentage of chemical class - Kruskal-Wallis with Dunn’s post-hoc test).

### Chemical analysis

Chemical analysis of *B. impatiens* DG secretion revealed a highly caste specific chemical composition that varies with life stage, reproductive, mating and social condition. The DG secretion varied significantly in the proportion of alkanes (Kruskal Wallis test, χ^2^ = 50.47, df = 3, p < 0.001), alkenes (Kruskal Wallis test, χ^2^ = 51.79, df = 3, p < 0.001), esters (Kruskal Wallis test, χ^2^ = 59.41, df = 3, p < 0.001), and terpenes (Kruskal Wallis test, χ^2^ = 60.84, df = 3, p < 0.001). Groups with activated ovaries had higher proportions of alkanes (QRW: 19 ± 0.4, gynes: 12.2 ± 1, QLW: 27.6 ± 1.6, queens: 2 ± 0.4; mean ± SE, Kruskal-Wallis with Dunn’s post hoc test, p < 0.05) and alkenes (QRW: 55 ± 1.5, gynes: 45.8 ± 1.1, QLW: 66.7 ± 1.8, queens: 67.1 ± 1.4; mean ± SE, Kruskal-Wallis with Dunn’s post hoc test, p < 0.05), while groups with inactivated ovaries, QRW and gynes, had a significantly higher proportion of esters than those with activated ovaries (QRW: 26.1 ± 1.7, gynes: 36.5 ± 1.8, QLW: 5.7 ± 0.8, queens: 2 ± 0.4; mean ± SE, Kruskal-Wallis with Dunn’s post hoc test, p < 0.05) (Fig 1b). Terpenes were present in queens and gynes, but not workers, and were higher in gynes compared to any other female group (QRW and QLW: not detected, gynes: 5.6 ± 0.4, queens: 2.07 ± 0.5; mean ± SE, Kruskal-Wallis with Dunn’s post hoc test, p < 0.05). NMDS analyses show that differences in the relative composition of DG compounds distinguish between groups based on caste (queens vs. workers), life stage, mating and reproductive status (queens vs. gynes) and social and reproductive condition (QRW vs. QLW) (Figure S1). PERMANOVA analyses of the relative composition of individual components of the DG secretion showed significant differences between all groups, regardless of chemical class (FDR correct p-values all < 0.01, Table S2).

### RNAseq results

Of the 9569 genes used in differential expression analyses, more than a third were differentially expressed between queens and gynes (1381 upregulated and 1271 downregulated). A similar number were differentially expressed between queen and worker castes (1405 upregulated and 1344 downregulated). In contrast, there were no significantly DEGs between the worker treatments, which also clustered together in a t-SNE plot (Fig 2). A heatmap of the 100 most DEGs (ranked by log-odds ratio) for each contrast shows striking differences between queens, gynes, and workers, but clustering of QRW and QLW treatment groups (Fig 3). Volcano plots show the large magnitude of expression differences between all treatment groups except between QRW and QLW (Fig S2). Interestingly, the number and magnitude of DEGs between queens and gynes were greater than between any queen-workers comparison, and varied up to a log fold change of 10 (Fig S2). An Upset plot (Fig 4) shows the overlap of DEGs between different contrasts, and identifies the number of genes which vary in response to caste (differentially expressed in queens and gynes vs. workers, 239 DEGs), reproduction (differentially expressed between groups with activated and inactivated ovaries (queens and QLW vs. gynes and QRW), 82 DEGs), and mating status (differentially expressed in mated queens vs. gynes and workers, 526 DEGs).

**Figure 2.**
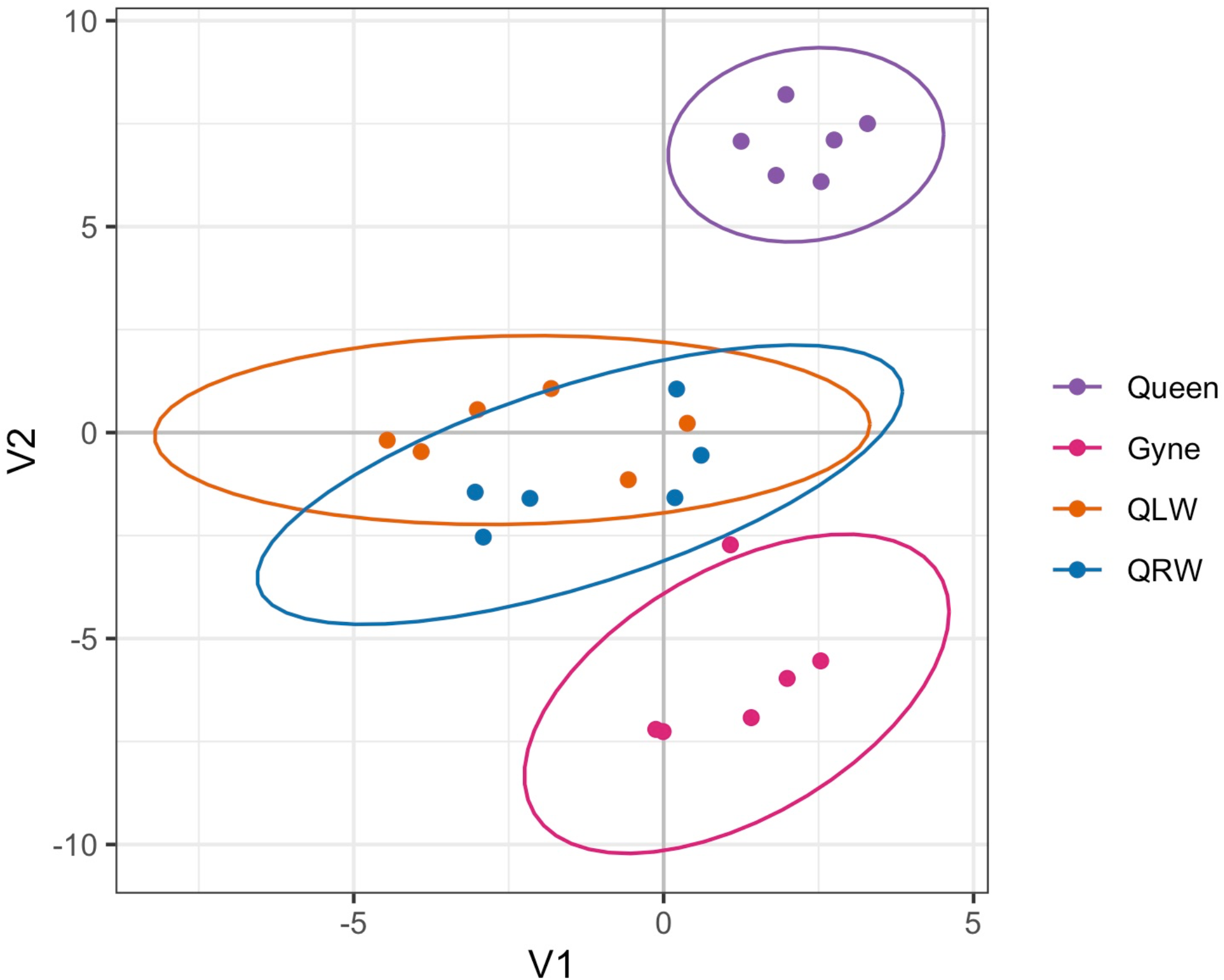
t-SNE plot showing clustering of Dufour’s gland RNAseq samples based on treatment group using all gene expression data.

**Figure 3.**
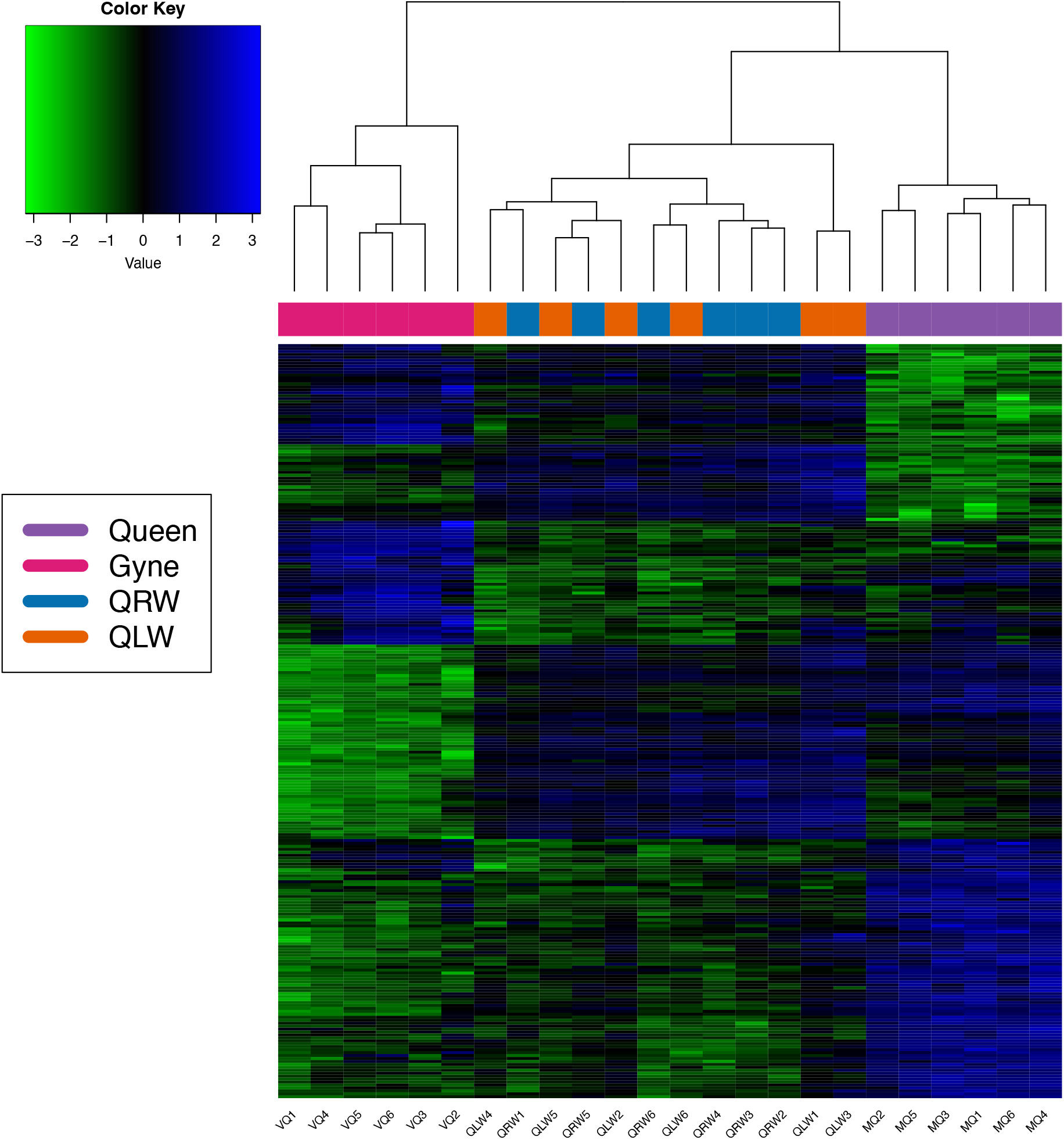
Heatmap showing the scaled (mean = 0, sd = 1), log2 expression values for a subset of the differentially expressed genes for each sample of the RNAseq experiment (6 samples per treatment and a total of 24 libraries). Rows are genes and columns are samples. The subset of genes was created by selecting the top 100 differentially expressed genes based on the highest absolute value of fold change for each contrast. This list produced 297 unique genes used in the heatmap. Colors at the tips of the dendrogram represent the treatment group of the samples.

**Figure 4.**
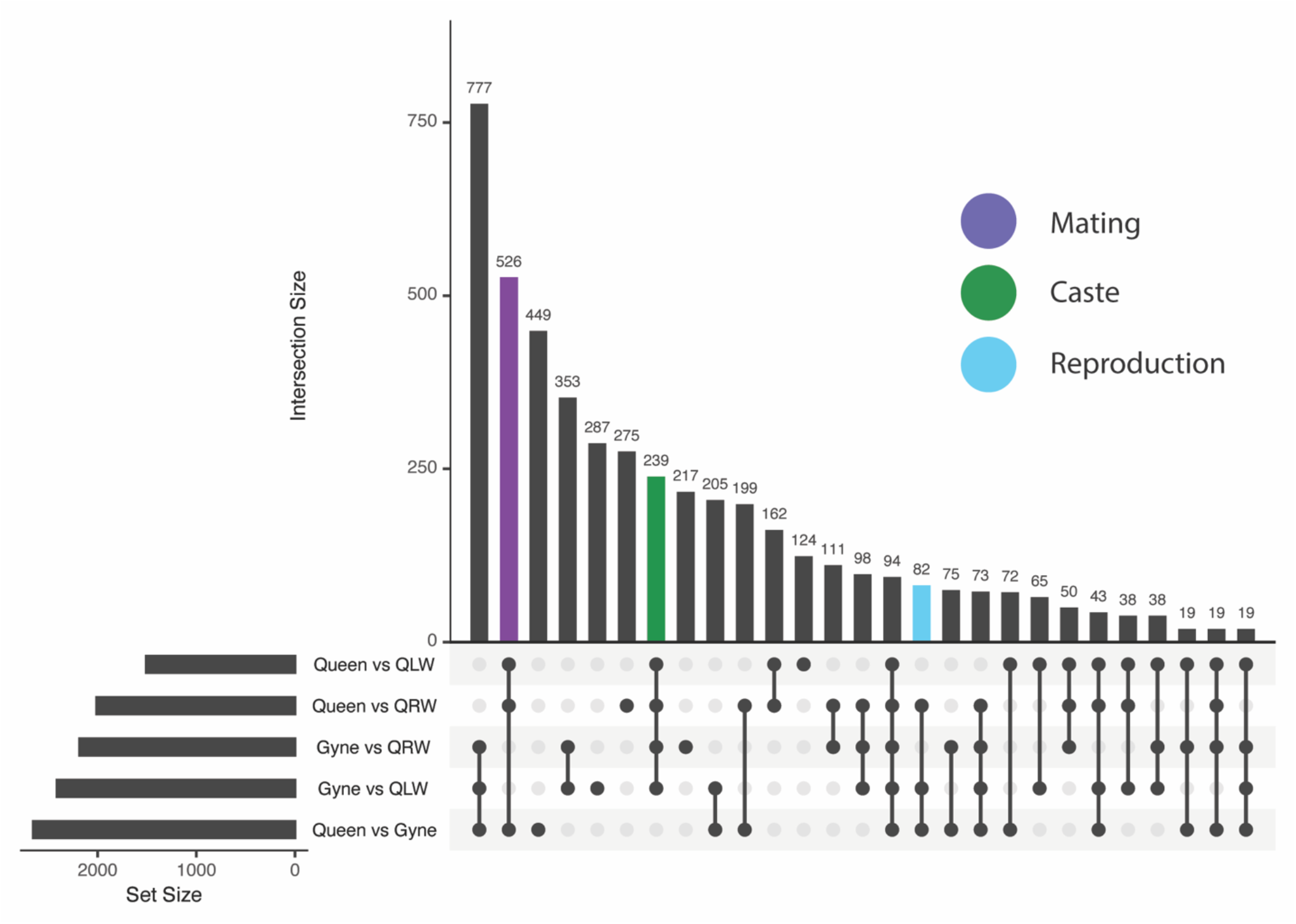
Upset plot showing the overlap of differentially expressed genes in the Dufour’s gland between all pairwise comparisons. Colored bars represent gene lists likely to be enriched for genes related to mating status (purple), caste (green), and reproductive state (blue), selected by identifying overlap with comparisons sharing a given trait. Pairwise comparisons were between treatment the treatment groups “queen” (mated, egg-laying queens > 2 months old), “gyne” (unmated, non-reproductive 6-day old gynes), “QRW” (7-day old queenright workers), and “QLW” (7-day old queenless workers).

To identify genes responsible for the differences in DG secretion between the treatment groups, we examined genes related to the fatty acid biosynthetic pathway which produce wax esters (produced by workers and gynes, proposed biosynthesis is provided in Fig 5), and the mevalonate/isoprenoid biosynthetic pathway which produces terpenes (produced mostly by gynes, but also by queens, Fig 6). Since QRW and QLW were indistinguishable in terms of gene expression, they were grouped together as one variable in the plots identifying genes involved in biosynthesis.

**Figure 5.**
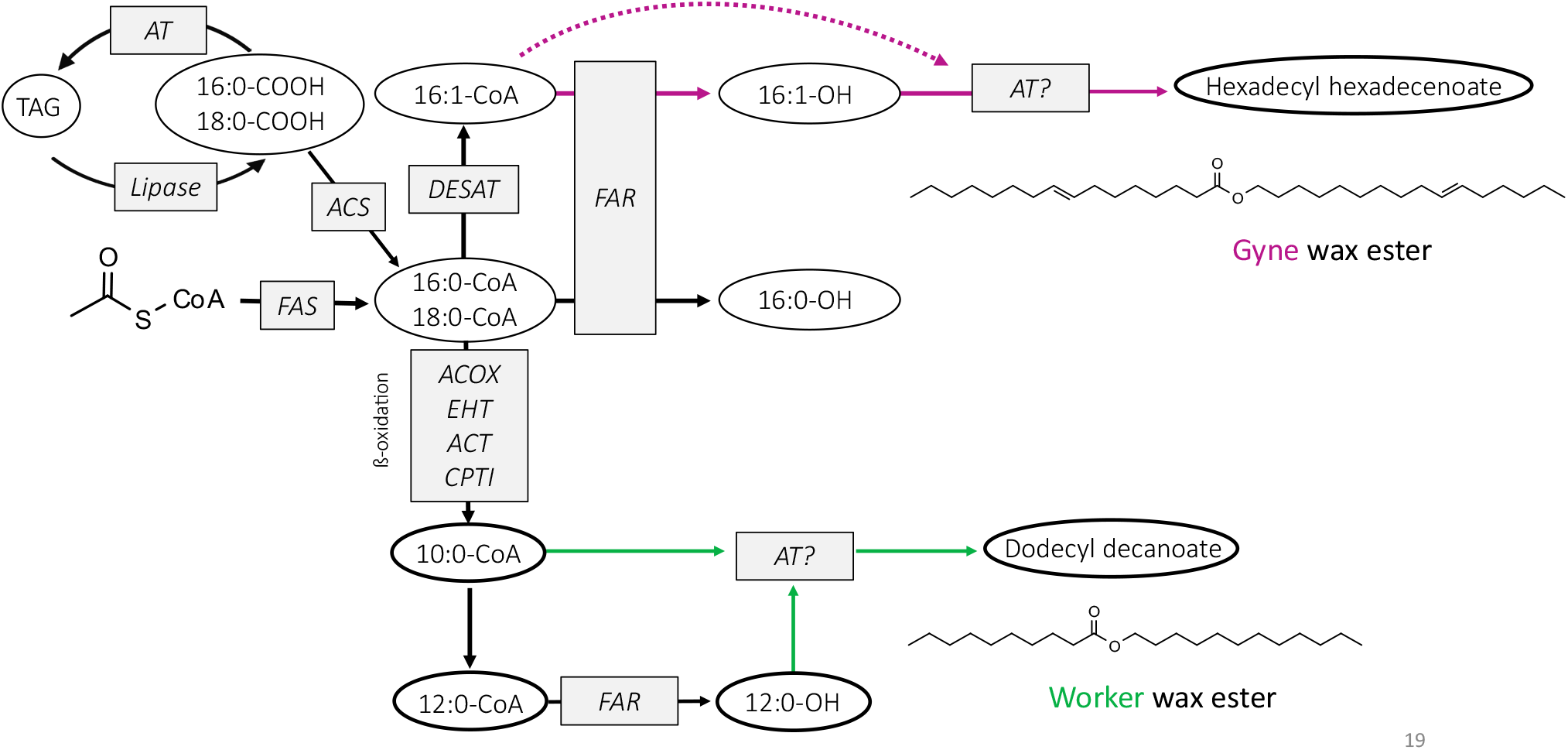
Proposed biosynthetic pathway of gyne-like or worker-like wax esters (figure adapted from Buček et al 2017). Enzymes abbreviations in the pathway are: *FAS* (fatty acid synthase), *ACOX* (acyl-CoA oxidases, *EHT* (enoyl-CoA hydratases), *ACT* (3-ketoacyl-CoA thiolases), *CPTI* (carnitine O-palmitoyltransferase 1, liver isoform), *DESAT* (fatty acyl desaturases), *FAR* (fatty acyl reductases), *AT* (acyltransferases), *ACS* (acyl-CoA synthases).

**Figure 6.**
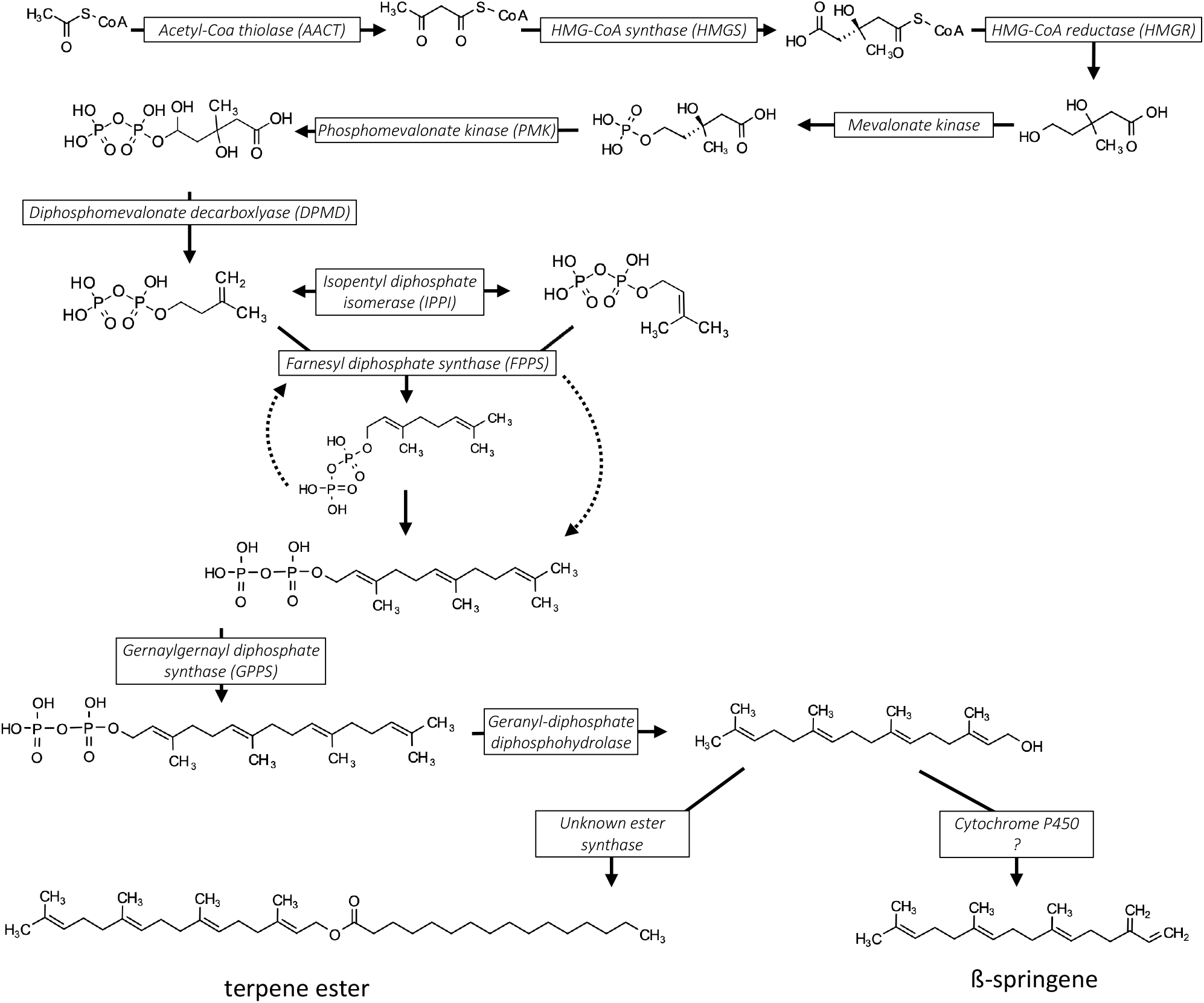
Proposed biosynthetic pathway of terpenes and terpene esters which are unique to the queen caste (adapted from Buček et al 2017). The figure depicts how geranylgeraniol could either be reduced to produce β-springene by an unknown enzyme or combined with a fatty acid to produce a terpene ester.

### Genes regulating wax esters in females

Genes related to fatty acid biosynthesis included enzymes involved in 5 discrete steps of wax ester biosynthesis. The first step, fatty acid synthesis, is responsible for the earliest steps in the construction of fatty acid chains from acetyl-CoA units (Fig 7A). Here, two DEGs involved in this process were recovered, while fatty acid synthase genes did not vary between treatments. We found 7 DE genes involved in β-oxidation, which shortens fatty acid chains by two carbons. These tended to be more expressed in workers and queens than gynes (Fig 7B). Fatty acyl reductases (FAR) produce fatty alcohols from fatty acyl-CoA and depending on the specific gene, varied between all treatment groups (Fig 7C). About half of the FAR genes (7 of 15) were exclusively upregulated in workers compared to queens and gynes. The remaining FAR genes varied in their expression patterns between groups with six being downregulated in workers compared to queens, gynes or both, and the two other being upregulated in both workers and queen compared to gynes (Fig 7C). Desaturases (DESAT) introduce double-bonds to fatty acyl-CoA and all three DE genes were highest in gynes, differing mostly between gynes vs. queens and workers, but not between queens and workers (Fig 7D). Finally, acyltransferases (mono and diacylglyceroltransferases MGAT/DGAT), synthesize mono, di, and triglycerides, in addition to a possible role as wax synthases and were most highly expressed in gynes (3 of 4 genes) (Fig 7E). Elongases lengthen the chain of fatty acyl-CoA beyond 16 or 18 carbons and tended to be more highly expressed in gynes than workers or queens (Fig S4). As the long chain wax esters differentiating the queen caste are comprised of acid and alcohol portions up to 18 carbons, these are more likely to be involved in cuticular hydrocarbon than wax ester biosynthesis.

**Figure 7.**
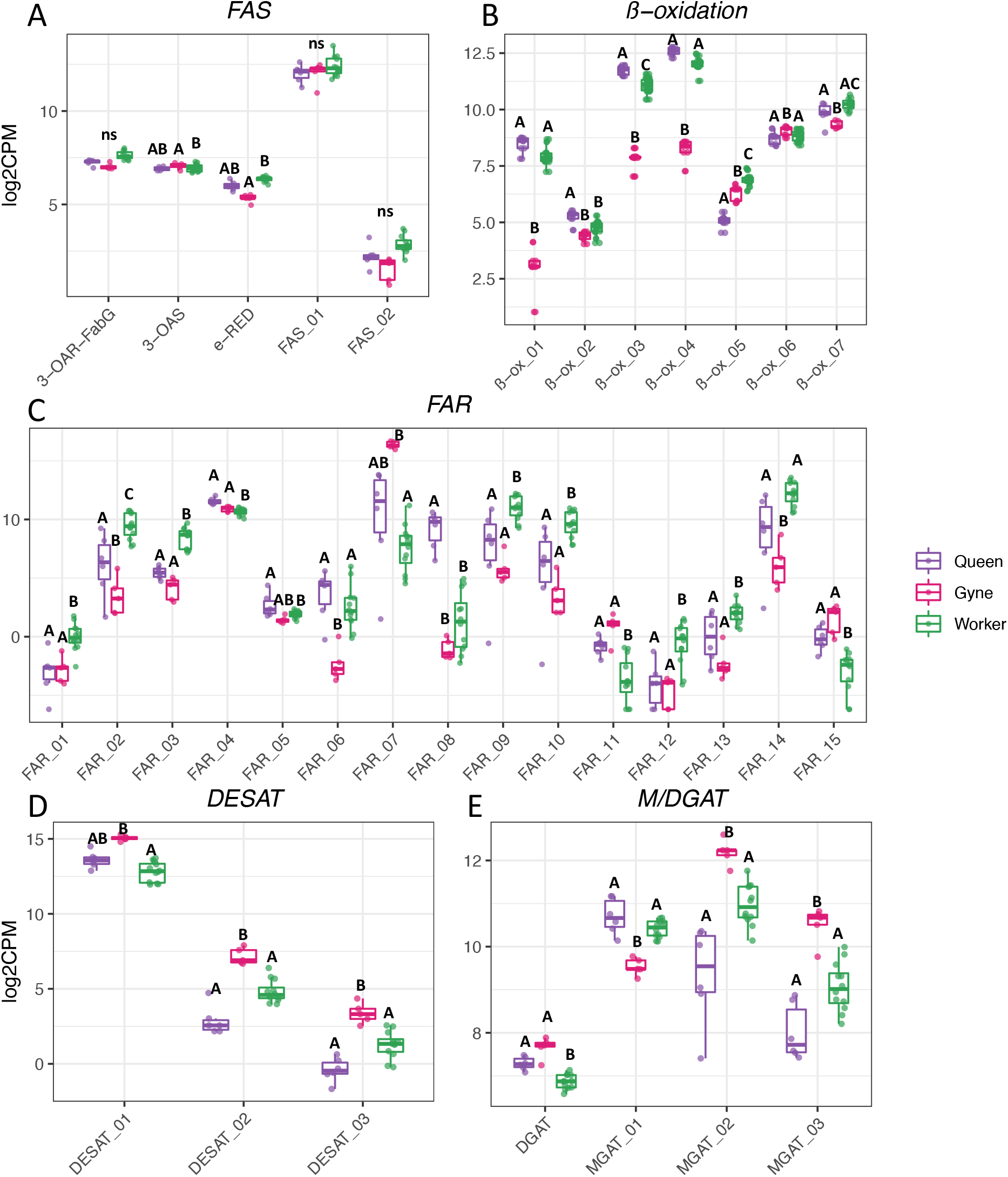
DEGs from RNAseq analyses involved in fatty acid biosynthesis. Gene IDs and annotations of labeled genes are given in Table S3. Data are presented as boxplots overlaid with each data point. Letters above columns denote statistical differences of pairwise contrasts using Benjamini and Hochberg adjusted p-values at α = 0.05.

### Genes regulating terpenes in females

Differentially expressed genes related to isoprenoid/mevalonate biosynthesis included genes involved early in the pathway and several genes downstream the pathway. The early genes (AACT, HMGS, HMGR, PMK) construct the precursors to all terpenes and terpenoids and function in the production of isopentyl diphosphate from acetyl-CoA, while downstream genes like farnesyl pyrophosphate synthase (FPPS) condenses isopentyl diphosphate (IPP) and dimethylallyl diphosphate (DMAPP) to make geranyl and subsequently farnesol diphosphate, the precursor of all future farnesyl compounds. Farnesol dehydrogenase genes oxidize farnesol to farnesal, for instance in the production of juvenile hormone that serves as a gonadotropin in female insects (Mayoral, Nouzova, Navare, & Noriega, 2009; Rivera-Perez, Nouzova, Lamboglia, & Noriega, 2014). Genes related to the isoprenoid/mevalonate pathway were among the most highly expressed genes we detected (Fig S3). Genes in the early isoprenoid/mevalonate pathway, including HMGR, the rate-limiting enzyme for the whole pathway, tended to be most highly expressed in gynes, which also produced the largest amounts of terpenes and terpene esters. The farnesol dehydrogenase genes were all most highly expressed in queens, while gynes and workers did not differ (Fig 8). Gene IDs and annotations corresponding to genes displayed in Figures 7 and 8 are given in Table S3, and statistical values of each group contrast given in Supplementary File S1.

**Figure 8.**
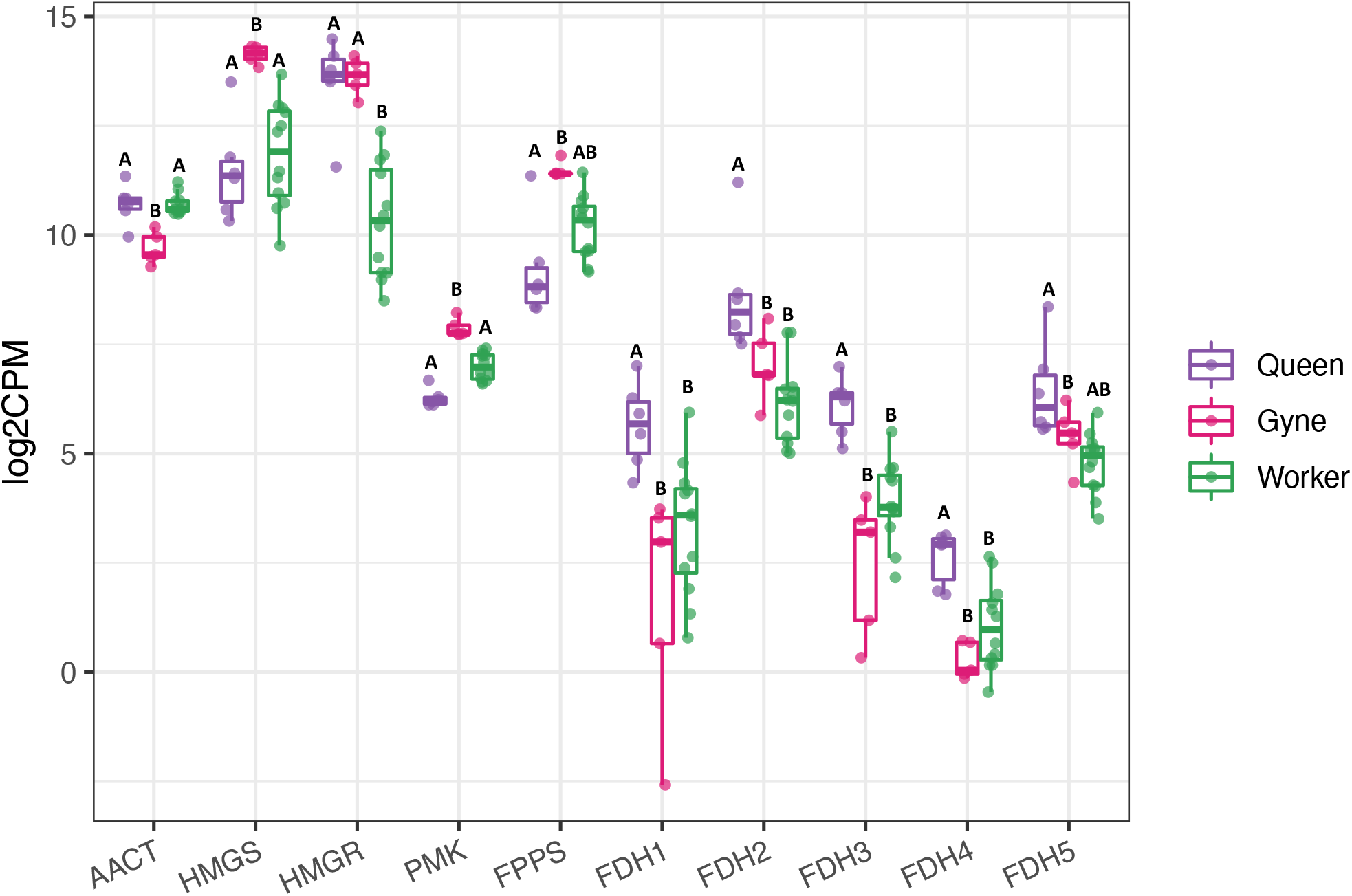
DEGs from RNAseq analyses in the isoprenoid/mevalonate biosynthetic pathway. Gene IDs and annotations of labeled genes are given in Table S3. Data are presented as boxplots overlaid with each data point. Letters above columns denote statistical differences of pairwise contrasts using Benjamini and Hochberg adjusted p-values at α = 0.05.

## Discussion

In the current study we examine the chemical composition and patterns of gene expression in the Dufour’s gland of *Bombus impatiens* with regard to caste, reproductive, and mating status. We focus on genes predicted to participate in the biosynthesis of esters, terpenes, and terpenoids that likely act as signals mediating social or mating behaviors. As predicted, we found that many of the enzymes involved in fatty acid and isoprenoid/mevalonate biosynthesis were significantly expressed and differentially regulated across castes, highlighting the role of the gland in semiochemical biosynthesis and providing many candidate genes for further investigation. Overall, differential expression analyses showed large differences between the DG of gynes and workers (∼25% of detected genes) and queens and workers (∼30% of detected genes), while QRW and QLW were indistinguishable.

Chemical analyses reveal the DG composition to be distinct among queens, gynes, QLW, and QRW in line with previous work in *B. impatiens* (Derstine et al., 2021). In gynes and queens, this is primarily explained by the production of longer chain wax esters (e.g. hexadecyl hexadecenoate) and terpene compounds (β-springene and isomers in addition to unidentified terpene esters that are absent in workers). However, also the relative amounts of various chain length alkanes and alkenes is specific to caste and reproductive state. Further differentiating the castes are a series of worker-specific dodecyl esters which are produced in larger amounts by queenright compared to queenless workers and are not produced by queens. These worker-specific esters are found in higher proportion in QRW with inactive ovaries (Derstine et al., 2021).

Linking chemical and gene expression phenotypes, we found many candidate genes likely involved in the biosynthesis of DG compounds. Among these were FARs and M/DGATs involved in wax ester biosynthesis, as well as high expression of genes in the isoprenoid/mevalonate pathway that leads to terpenes. Fatty-acyl desaturases, a class of enzymes in the fatty acid biosynthesis pathway which introduce double bonds, were all upregulated in gynes compared to queens and workers. This matches the chemistry data from the Dufour’s gland showing that gynes produce the largest amounts of both unsaturated hydrocarbons and wax esters with unsaturated alcohol or acid moieties. Similarly, the acid moiety of most of the long chain wax esters found in gynes contains unsaturations. Worker wax esters are further differentiated from queens and gynes by decreased length, which could be explained by an upregulated β-oxidation chain-shortening process. This iterative reduction in fatty acid chain length by two carbons occurs in both mitochondria and peroxisomes, and β-oxidation genes involved in both locations are differentially expressed in the Dufour’s gland (Demarquoy & Le Borgne, 2015). However, the peroxisomal genes (β-ox_01-04, Fig 7b) are more involved in biosynthetic as opposed to catabolic processes, and accordingly, show much larger fold change differences in workers and queens compared to gynes.

Interestingly, we find that caste is much more explanatory of chemical phenotypes than reproductive state, a finding highlighted by the lack of DEGs between QRW and QLW which vary drastically in ovarian activation. Gene expression data did not explain the quantitative differences in the DG composition of worker groups, or more generally reflect any differences related to reproductive state, although the DG opens into the oviduct and is associated with the reproductive system of females. It is possible that differences in DG composition between QRW and QLW could result from the differential secretion of certain compounds, while the production remains the same. For instance, QLW which have a lower percentage of esters could be secreting them at a higher rate than QRW in line with a reproductive signaling function. Alternatively, regulation of these compounds could happen at the protein instead of the transcript level, which we would not detect with RNA sequencing.

Wax esters in the Dufour’s gland could be biosynthesized via two different routes, either de novo, or from fatty acids transported from outside the gland. In the first option, fatty acids synthesized outside the gland are stored as triacylglycerol (TAG) in fat body tissue. Specific lipases act on TAGs to produce diacylglycerol (DAG), which is bound to the transport protein lipophorin and transported to various tissues (Majerowicz & Gondim, 2013; Schal, Sevala, & Cardé, 1998). Once inside the cell, lipases release fatty acids from DAG where they are converted to their final form by the necessary array of enzymes. In the de novo route, fatty acids are constructed from acetate units inside the cell, where the same array of enzymes would convert them to their final form. Thus, expression of genes early in the fatty acid biosynthetic pathway provide evidence for de novo biosynthesis. This may be indicated by the expression of fatty acid synthase (FAS), which combines acetate units to produce 16 and 18 carbon fatty acyl-CoA molecules. While not differentially expressed between the groups, FAS has relatively high expression compared to many genes found in the study. Furthermore, the expression of numerous elongases which extend fatty acyl-CoA molecules beyond 16 and 18 carbons is consistent with de novo hydrocarbon synthesis. Previous work in honey bees found that the hydrocarbons are likely transported into the DG while the esters are synthesized de novo (Gozansky, Soroker, & Hefetz, 1997). This has yet to be established in *Bombus*.

Similarly, early genes in the terpene pathway, including AACT, HMGS, and HMGR, the rate limiting enzyme of the isoprenoid pathway (Friesen & Rodwell, 2004), were highly expressed relative to other genes we detected, providing strong evidence for de novo terpene synthesis. The pathways first steps are shared by almost all organisms and result in the production of isopentyl diphosphate (IPP) and dimethylallyl diphosphate (DMAPP) from acetyl-CoA (Miziorko, 2011). These two molecules act as the substrates for a variety of enzymes, which can be functionalized and combined in various iterations to produce the plethora of mono-, sesqui-, and di-terpenes and terpenoids (Darragh et al., 2021; Miziorko, 2011; Morgan, 2010), including those commonly found in various bumble bee exocrine glands, such as β-springene, dihydrofarnesol, and terpene esters such as the fatty acid esters of geranylcitronellol or farnesol (Derstine et al., 2021; Abraham Hefetz, Taghizadehr, & Francke, 1996; Orlova, Villar, Hefetz, Millar, & Amsalem, 2021). Overall, the high levels of expression and numerous DEGs in fatty acid and isoprenoid biosynthetic pathways are consistent with de novo pheromone synthesis of esters and terpenes in the Dufour’s gland.

Throughout nature, wax esters are biosynthesized by enzymes in the acyltransferase family, but no specific wax synthases have yet been identified in insects. In mice, wax esters are biosynthesized by two acyltransferases, a multifunctional *DGAT1*, which has moderate ability to produce wax esters, primarily catalyzing the production of triacylglycerols, and a “dedicated” wax synthase, which produces wax monoesters but has little ability to catalyze other types of esters (Cheng & Russell, 2004; Yen et al., 2005). Plants and bacteria have either a specific wax synthase enzyme, or multifunctional acyltransferases such as DGATs which both catalyze fatty acid ester formation and triacylglycerols. In the honey bee *A. mellifera*, the monoesters found in beeswax are typically composed of C16-C20 acids and C24-32 or larger alcohols, and are synthesized by an acyltransferase enzyme, as shown by studies using radio-labeled precursors (Blomquist & Ries, 1979). A candidate wax synthase gene in the acyltransferase family was identified in the white wax scale *Ericerus pela* by comparing gene expression levels in instars, but has yet to be functionally characterized (Yang et al., 2012).

Terpene biosynthesis is famously diverse, and as a consequence we would predict genetic regulation of this pathway to be less specific than in wax ester biosynthesis. To narrow down the focus, we selected a specific tissue (e.g. the Dufour’s gland is not predicted to participate in the biosynthesis of other common insect terpenes, like corpora allata is in producing juvenile hormone), and focused on the rate limiting enzymes. Two genes responsible for terpene synthesis, HMGS and HMGR, were DE in queens and/or gynes compared to workers, potentially explaining the presence of terpenes and terpenes esters that are absent in workers. Data from other species of *Bombus* male show transcript abundances of some of the same DE isoprenoid biosynthesis genes (AACT, HMGR, FFPS) correlated with observed differences in male pheromone components of *B. lucorum* which contains primarily fatty-acid derived compounds (ethyl tetradecenoate and ethyl dodecanoate), and *B. terrestris* which contains primarily isoprenoid compounds (E-2,3-dihydrofarnesol, geranylcitronellol, and 2,3-dihydrofarnesal) in addition to some fatty acid derived compounds (ethyl dodecanoate, octadeca-9,12,15-trienol, and hexadecan-1-ol) (Prchalova et al., 2016). However, while gynes clearly have the largest amount and proportion of terpene compounds, many of the genes in the pathway often have as low or lower expression in workers, which produce none. HMGR is the rate-limiting enzyme in this pathway, and is under extensive regulation at the level of expression, translation, and post-translation (Friesen & Rodwell, 2004). This coupled with the diverse downstream products of terpene synthesis help to explain the expression seen in worker groups which do not produce terpenes.

Biosynthetic studies of other organisms with terpene pheromones also found expression of early mevalonate pathway genes in pheromone producing tissues, coupled with highly specific but often unknown downstream enzymes such as terpene synthases. This is the case in the psychodid fly *Lutzomyia longipalpis*, which produces a blend of homosesquiterpenes that function as a male sex pheromone (González-Caballero et al., 2014; González-Caballero, Valenzuela, Ribeiro, Cuervo, & Brazil, 2013), and in the terpene based male labial gland components of *Bombus terrestris* (Bucek et al., 2016; Prchalova et al., 2016). Until recently, no terpene synthases had been identified in insects, but it has now been shown in several lineages that insects have co-opted known genes in the isoprenoid pathway. For instance, enzymes with terpene synthase activity in flea beetles (Beran, Rahfeld, et al., 2016), stink bugs (Lancaster et al., 2018; Lancaster et al., 2019) and a *Heliconius* butterfly (Darragh et al., 2021) derive from an isoprenyl diphosphate synthase (IDS).

Understanding the biosynthesis and genetic regulation of insect pheromones allows the study of their evolution, which can inform important processes like speciation and the evolution of social behavior. For instance, allelic variation in the reductase pgFAR explains the different pheromone blends produced by two strains of European corn borer moths, which leads to reproductive isolation and probable early steps of speciation in the field (Lassance et al., 2010). Similarly, the distinct pheromone blends of different genera of leafroller moths are attributed to differential regulation of the same desaturase gene (Albre et al., 2012). In social insects, reproductive division of labor is regulated in part by chemical communication, with different castes signaling fertility or producing pheromones which regulate ovarian activation or egg-laying (A. Hefetz, 2019; Y. Le Conte & Hefetz, 2008). In many species, these females have the same genotype, and thus the different pheromones or pheromone blends they produce are likely the result of differential gene expression. In honey bees, queens regulate worker reproduction with a multi-component pheromone blend from the mandibular gland, the major components of which are 9-oxo-2-decenoic acid (9-ODA) and 9-hydroxy-2-decenoic acid (9-HDA) (Hoover et al., 2003; Slessor et al., 1990), while the major product in workers is 10-hydroxy-2-decenoic acid (10-HDA) (E Plettner, Sutherland, Slessor, & Winston, 1995; Slessor, Kaminski, King, Borden, & Winston, 1988). Research to understand the caste-specific biosynthesis of these compounds identified the hydroxylation step as the probable divergence point between queen and worker chemical phenotypes, and posited that the key enzymes were CYP450 hydrolases with different substrate affinities (Erika Plettner et al., 1998). This was confirmed in later work that identified differential expression between CYP4AA1 (worker hydroxylation pattern) and CYP18A1 (queen hydroxylation pattern) as producing the observed patterns of caste-specific hydroxy acid biosynthesis (O. Malka, Karunker, Yeheskel, Morin, & Hefetz, 2009). Furthermore, this expression can be plastic, with queenless workers able to switch pathways and produce more queenlike secretions (Crewe & Velthuis, 1980; Osnat Malka, Shnieor, Hefetz, & Katzav-Gozansky, 2006; E Plettner, Slessor, Winston, Robinson, & Page, 1993). These insights have helped explain biological phenomena such as the interesting case of the honey bee subspecies *A. m. capensis*, whose workers are capable of becoming social parasites in part by upregulating the queen specific hydroxylation pathway to produce higher amounts 9-ODA and 9-HDA (F. N. Mumoki, Yusuf, Pirk, & Crewe, 2019; Fiona N Mumoki, Yusuf, Pirk, & Crewe, 2021). Additionally, the evolution of queen pheromones is thought to be an important step in the evolution of eusociality, and understanding the genes involved is necessary for comparative studies with solitary ancestors that seek to understand the evolutionary origin of pheromones regulating reproduction (Treanore, Derstine, & Amsalem, 2021).

Finally, matching gene expression to chemical phenotypes is complicated by the multifunctional nature of many of the enzymes involved, and the limited functional information available for the genes of interest. For example, the synthesis of dodecyl esters produced by *B. impatiens* workers are likely to involve activity by FAR and DGAT in the final synthetic step. However, DGAT genes likely catalyze not only wax ester formation but also triacylglycerides. Additionally, FAR enzymes play a role in the reduction of fatty-acyl molecules, a process taking place in all castes, regardless of caste-specific pheromone production, and little is known about the substrate specificity of the numerous FAR enzymes. Furthermore, it is unknown if similar signals produced by different castes (e.g., shorter chain esters in workers and long chain/terpen esters in gynes) are synthesized by the same enzymes. Therefore, multifunctional enzymes pose a challenge and do not necessarily result in direct matches between chemical and gene expression phenotypes.

Overall, our data provide ample opportunities to further the understanding of signal evolution and their genetic regulation in social insects with functional studies targeting specific genes, and experiments designed to assess the specific reactions and substrate specificity catalyzed by the many candidate enzymes described. As genes involved in the production of social signals are more likely to have undergone recent selection, it would also be fruitful to compare rates of evolution and gather evidence about the type of selection experienced by these genes during the transition from their solitary past.

## Acknowledgements

This work was funded by NSF-CAREER IOS-1942127 to EA.

## Author contribution

ND and EA designed the experiments. ND and GV performed the experiments. ND and DG conducted data and statistical analyses. ND and EA wrote the manuscript. All authors provided feedback about the final draft.

## Data Accessibility and Benefit-Sharing Statement

Raw sequence reads and metadata will be deposited at NCBI upon manuscript acceptance. Chemical data from the analysis of the Dufour’s gland secretion and terminal oocyte measurements of bees used in the study are available from the authors upon request.

