## Supplementary Material for "Differential gene expression underlying the biosynthesis of Dufour’s gland signals in *Bombus impatiens*"

RNAseq Supplementary Info

Figure S1. NMDS plots based on the Bray-Curtis dissimilarities of relative peak areas of Dufour's gland compounds in 4 treatment groups (queens, gynes, queenless workers – QLW, and queenright workers – QRW). Plots were created using only ester, only alkanes, only alkenes, or all compounds. PERMANOVA tests using Bray-Curtis dissimilarity matrices showed significant differences in all treatment categories when using any chemical class.

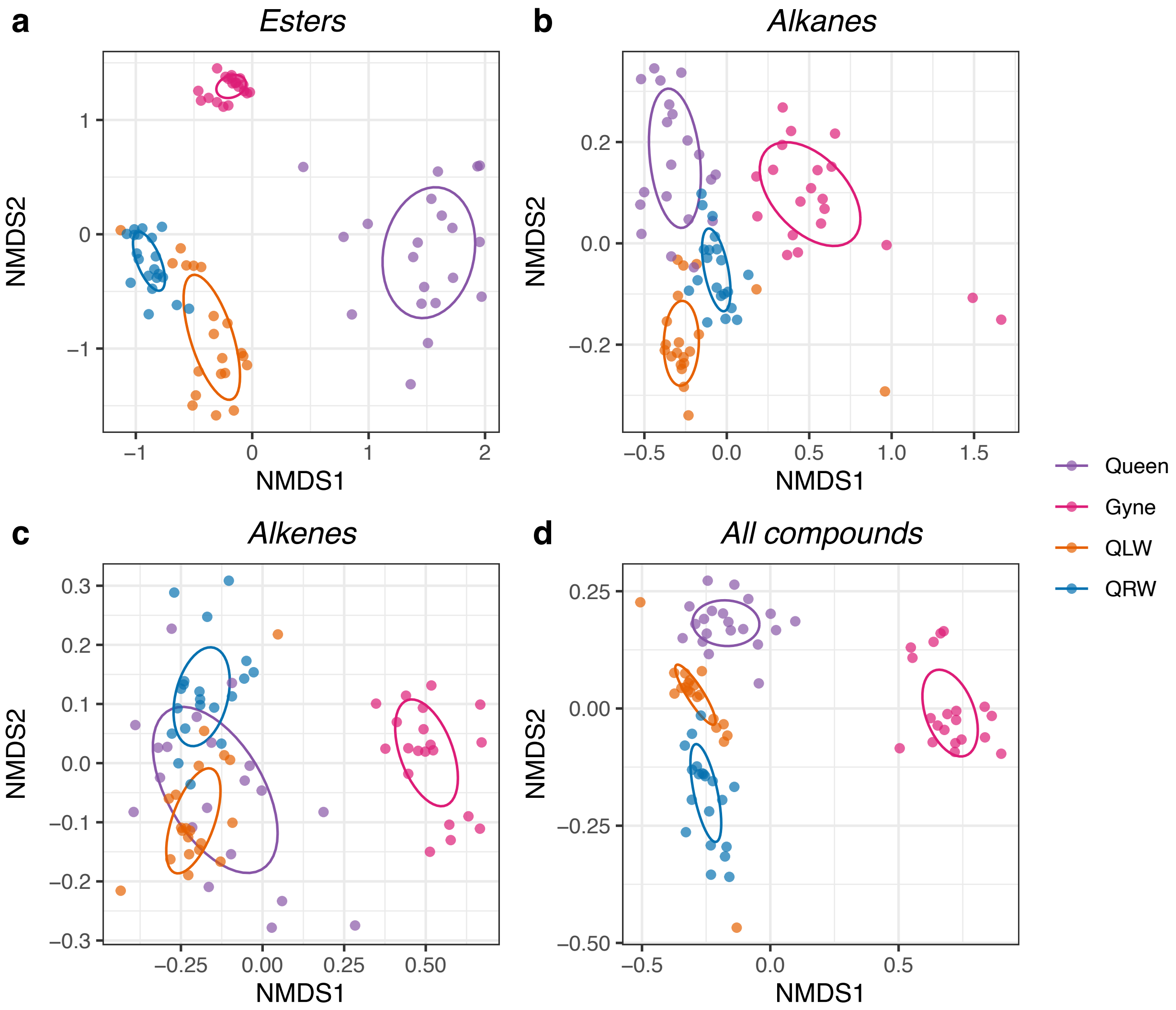

Figure S2 – Volcano plots showing the log2 fold change (x-axis) against the log10 adjusted p-values for genes within each specific contrast. Red points have a significant p-value and a fold change > 1.5, blue points are significant but below the fold-change cut-off. Green and gray points are not significant.

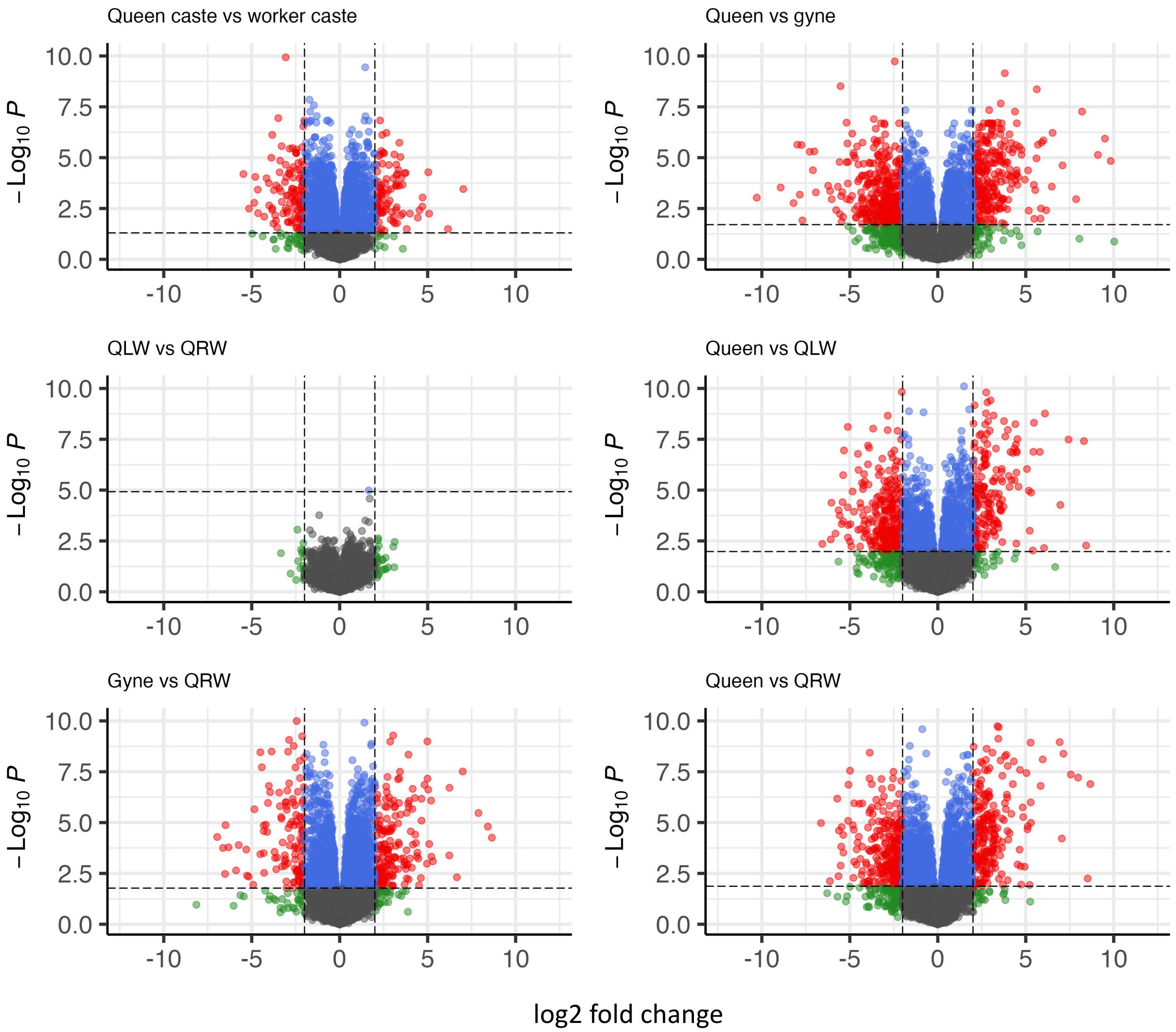

Figure S3 – Mean TPM (transcripts per million mapped reads) per treatment group, ranked by decreasing abundance. The TPM calculation normalizes for sequence length and library size. The top 1000 genes are displayed.

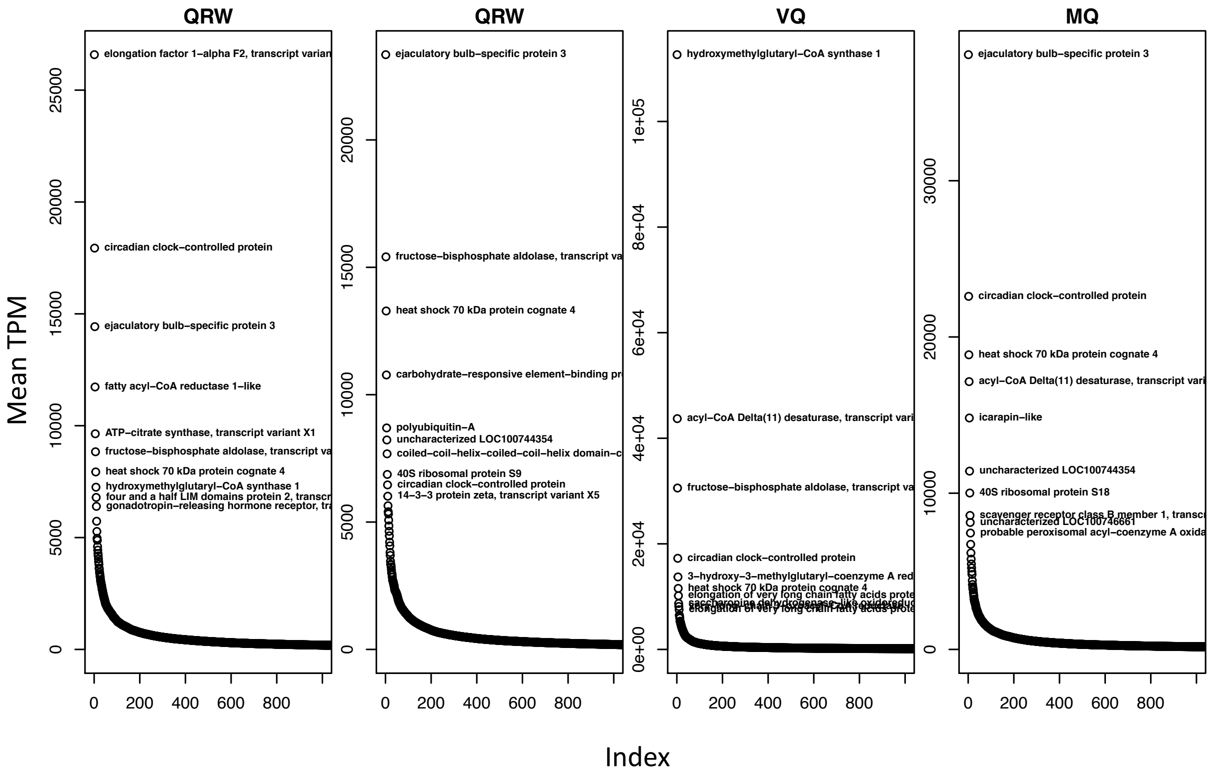

Figure S4 - Elongases potentially involved in CHC synthesis, lengthening fatty acyl-CoA molecules beyond 18 carbons.

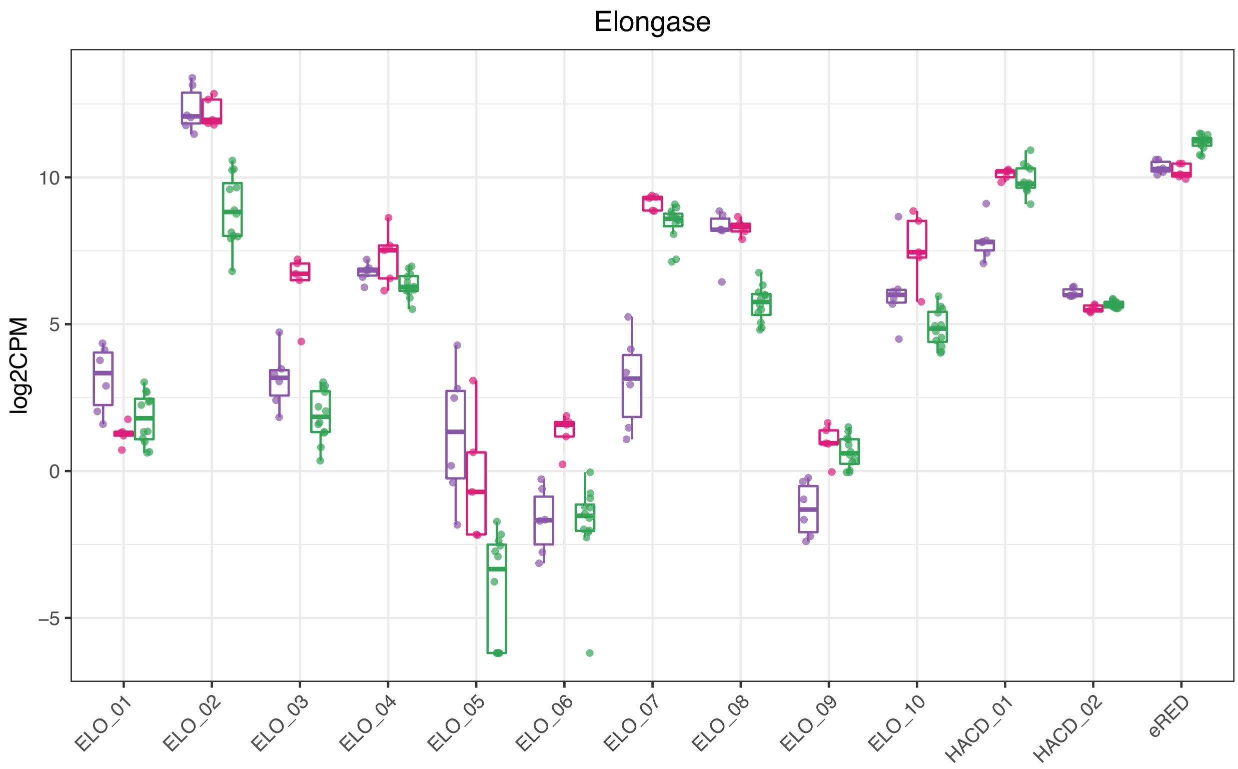

Table S1. Information for RNAseq samples, including the number of glands per pool, colony, age, and purity.

| Colony | Treatment | Tissue pool | Oocyte size mm (mean ± SE,) | caste | age | ID |
|---|---|---|---|---|---|---|
| 47 | Gyne | 3 | 0.21 ± 0.3 | queen | 3-4 days | 5A |
| 56 | Queen | 1 | 3.26 | queen | > 2 months | 13A |
| C1 | Queen | 1 | 3.50 | queen | > 2 months | 15A |
| C38 | Queen | 1 | 3.36 | queen | > 2 months | 18A |
| 60 | QLW | 10 | 2.30 ± 0.19 | worker | 7 days | 21A |
| 59 | QLW | 5 | 2.44 ± 0.07 | worker | 7 days | 23A |
| 58 | QLW | 5 | 2.35 ± 0.11 | worker | 7 days | 24A |
| 59 | QRW | 10 | 0.32 ± 0.02 | worker | 7 days | 31A |
| 57 | QRW | 10 | 0.24 ± 0.03 | worker | 7 days | 33A |
| 63 | QRW | 9 | 0.21 ±0.02 | worker | 7 days | 39A |
| 66 | QRW | 9 | 0.77 ± 0.22 | worker | 7 days | 41A |
| 66 | QLW | 9 | 2.77 ± 0.06 | worker | 7 days | 42A |
| 62 | Gyne | 7 | 0.23 ± 0.03 | queen | 3-4 days | 44A |
| 63 | Gyne | 5 | 0.17 ± 0.04 | queen | 3-4 days | 45A |
| 65 | Gyne | 7 | 0.24 ± 0.03 | queen | 3-4 days | 46A |
| 66 | Gyne | 5 | 0.20 ± 0.03 | queen | 3-4 days | 47A |
| 64 | Gyne | 6 | 0.16 ± 0.02 | queen | 3-4 days | 48A |
| 66 | Queen | 1 | 3.28 | queen | > 2 months | 50A |
| 65 | Queen | 1 | 3.35 | queen | > 2 months | 51A |
| 64 | Queen | 1 | 3.32 | queen | > 2 months | 52A |
| 72 | QRW | 10 | 0.34 ± 0.06 | worker | 7 days | 53A |
| 72 | QLW | 10 | 2.20 ± 0.31 | worker | 7 days | 55A |
| 73 | QRW | 10 | 0.35 ± 0.06 | worker | 7 days | 57A |
| 73 | QLW | 10 | 2.79 ± 0.14 | worker | 7 days | 60A |

Table S2 – FDR corrected p-values of pairwise contrasts following PERMANOVA analysis comparing the relative composition of Dufour's gland compounds between queen, gyne, queenright workers, and queenless workers.

| | Queen - gyne | Queen - QR | Queen - QL | Gyne - QR | Gyne - QL | QR - QL |
|---|---|---|---|---|---|---|
| All compounds | 0.001 | 0.001 | 0.001 | 0.001 | 0.001 | 0.001 |
| Esters | 0.001 | 0.001 | 0.001 | 0.001 | 0.001 | 0.001 |
| Alkanes | 0.001 | 0.001 | 0.001 | 0.001 | 0.001 | 0.001 |
| Alkenes | 0.0012 | 0.0012 | 0.002 | 0.0012 | 0.0012 | 0.0012 |

Table S3 – Gene IDs and annotations of genes from Figure 8.

| gene_id | name | label |
|---|---|---|
| LOC100741857 | elongation of very long chain fatty acids protein 6 | ELO_01 |
| LOC100746103 | elongation of very long chain fatty acids protein 6 | ELO_02 |
| LOC100744329 | elongation of very long chain fatty acids protein AAEL008004 | ELO_03 |
| LOC100742554 | elongation of very long chain fatty acids protein AAEL008004-like | ELO_04 |
| LOC100743038 | elongation of very long chain fatty acids protein 6-like | ELO_05 |
| LOC100743508 | elongation of very long chain fatty acids protein AAEL008004 | ELO_06 |
| LOC100749115 | elongation of very long chain fatty acids protein 6-like | ELO_07 |
| LOC100749122 | elongation of very long chain fatty acids protein 6-like | ELO_08 |
| LOC100749526 | elongation of very long chain fatty acids protein 1 | ELO_09 |
| LOC100748215 | elongation of very long chain fatty acids protein 6 | ELO_10 |
| LOC100744206 | very-long-chain (3R)-3-hydroxyacyl-CoA dehydratase hpo-8 | HACD_01 |
| LOC100744962 | very-long-chain (3R)-3-hydroxyacyl-CoA dehydratase | HACD_02 |
| LOC100747299 | very-long-chain enoyl-CoA reductase | Enoyl_CoA_RED |
| LOC105680567 | 3-oxoacyl-[acyl-carrier-protein] reductase FabG | 3-OAR-FabG |
| LOC100747632 | enoyl-[acyl-carrier-protein] reductase | e-RED |
| LOC100742392 | 3-oxoacyl-[acyl-carrier-protein] synthase, mitochondrial | 3-OAS |
| LOC100742442 | fatty acid synthase | FAS_01 |
| LOC100742781 | fatty acid synthase | FAS_02 |
| LOC112213796 | peroxisomal acyl-coenzyme A oxidase 1-like | β-ox_01 |
| LOC100742496 | peroxisomal acyl-coenzyme A oxidase 3-like | β-ox_02 |
| LOC105681545 | probable peroxisomal acyl-coenzyme A oxidase 1 | β-ox_03 |
| LOC100747467 | probable peroxisomal acyl-coenzyme A oxidase 1 | β-ox_04 |
| LOC100749552 | short/branched chain specific acyl-CoA dehydrogenase, mitochondrial | β-ox_05 |
| LOC100746130 | probable enoyl-CoA hydratase, mitochondrial | β-ox_06 |
| LOC100749508 | carnitine O-palmitoyltransferase 1, liver isoform | β-ox_07 |
| LOC100744570 | putative fatty acyl-CoA reductase CG5065 | FAR_01 |
| LOC100746315 | putative fatty acyl-CoA reductase CG5065 | FAR_02 |
| LOC100746398 | putative fatty acyl-CoA reductase CG5065 | FAR_03 |
| LOC100740585 | putative fatty acyl-CoA reductase CG5065 | FAR_04 |
| LOC100746762 | putative fatty acyl-CoA reductase CG5065 | FAR_05 |
| LOC100745716 | putative fatty acyl-CoA reductase CG5065 | FAR_06 |

| | | |
|---|---|---|
| LOC100741032 | fatty acyl-CoA reductase wat | FAR_07 |
| LOC100750141 | putative fatty acyl-CoA reductase CG5065 | FAR_08 |
| LOC105680764 | putative fatty acyl-CoA reductase CG5065 | FAR_09 |
| LOC105681681 | fatty acyl-CoA reductase 1-like | FAR_10 |
| LOC100747705 | fatty acyl-CoA reductase 1-like | FAR_11 |
| LOC112214005 | putative fatty acyl-CoA reductase CG5065 | FAR_12 |
| LOC112214031 | fatty acyl-CoA reductase 1-like | FAR_13 |
| LOC100749776 | fatty acyl-CoA reductase 1-like | FAR_14 |
| LOC112214034 | putative fatty acyl-CoA reductase CG5065 | FAR_15 |
| LOC100746868 | acyl-CoA Delta(11) desaturase | DESAT_01 |
| LOC100746987 | acyl-CoA Delta(11) desaturase | DESAT_02 |
| LOC100749772 | acyl-CoA Delta(11) desaturase | DESAT_03 |
| LOC100746353 | 2-acylglycerol O-acyltransferase 1 | MGAT_01 |
| LOC100741228 | 2-acylglycerol O-acyltransferase 2-A-like | MGAT_02 |
| LOC100748623 | 2-acylglycerol O-acyltransferase 2-like | MGAT_03 |
| LOC100744121 | diacylglycerol O-acyltransferase 1 | DGAT |
| LOC100747910 | acetyl-CoA acetyltransferase, cytosolic | AACT |
| LOC100742345 | hydroxymethylglutaryl-CoA synthase 1 | HMGS |
| LOC100746066 | 3-hydroxy-3-methylglutaryl-coenzyme A reductase | HMGR |
| LOC100743984 | phosphomevalonate kinase | PMK |
| LOC100749317 | farnesyl pyrophosphate synthase | FPPS |
| LOC100747302 | farnesol dehydrogenase | FDH1 |
| LOC100742025 | farnesol dehydrogenase | FDH2 |
| LOC100747182 | farnesol dehydrogenase-like | FDH3 |
| LOC100743402 | farnesol dehydrogenase | FDH4 |
| LOC100741907 | farnesol dehydrogenase | FDH5 |
